# Quantitative STED microscopy with DNA-fluorophore labels

**DOI:** 10.64898/2025.12.05.692534

**Authors:** Laurell F. Kessler, Yunqing Li, Ashwin Balakrishnan, Mike Heilemann

## Abstract

Stimulated emission depletion (STED) microscopy enables super-resolution imaging of complex biological samples in 3D, in large volumes, and live. However, molecular quantification with STED has remained underexplored. Here, we present a straightforward approach for quantitative STED that enables molecule counting. For this purpose, we designed DNA-fluorophore labels that enable signal amplification and allow for reliable intensity-based quantitative imaging. We demonstrate accurate molecule counting on DNA origami. Furthermore, we visualized and quantified EGF receptor monomers and dimers in cells. In summary, we introduce a robust, fast, and easy-to-implement tool for quantitative STED microscopy with single-protein resolution.

## Introduction

Super-resolution microscopy (SRM) has revolutionized our understanding of cell biology ^1^. Stimulated emission depletion (STED) microscopy ^2^ is one of the major technologies in that field, which enables 3D, live-cell, multi-target super-resolution imaging, and has been applied in various cell biological systems across scales ^3,4^. Exciting recent developments, among others, are the integration of smart imaging workflows to detect rare events in living cells ^5^ or rare structures in high-throughput 3D imaging ^6^. Live-cell microscopy has been boosted by neural-network-assisted image reconstruction, enabling fast and long-term STED microscopy in living cells and 3D ^7,8^.

However, the extraction of quantitative information on the composition of nanoscale molecular assemblies, for example, membrane receptor clusters, remains underexplored for STED microscopy. Such questions are often addressed with quantitative single-molecule localization microscopy (SMLM) ^9–11^ or through direct visualization with true molecular resolution using MINimal fluorescence photon FLUXes (MINFLUX) ^12,13^. While these approaches have demonstrated impressive results, they either demand for a complex analysis, are restricted in the 3D sample volume that can be covered, or request advanced instrumentation and user experience. In contrast, STED microscopy is based on confocal scanning, and as such can cover large volumes in 3D and different types of samples. In addition, STED microscopy directly delivers an image without the need for computational image reconstruction steps, making it user-friendly. One reason that so far challenged quantitative analysis of nanoscale protein clusters is that an intensity analysis of single emitters in STED images might exhibit varying brightness across the sample. An elegant approach that mitigated this effect and enabled quantitative STED microscopy exploited the simultaneous photon arrival times in a time-correlated single-photon counting (TCSPC) imaging experiment, and reported molecular counting in super-resolved molecular clusters ^14^. However, this approach is experimentally demanding and requires complex instrumentation. A simple method for quantitative STED microscopy that is applicable to small molecular clusters is still missing, yet such a method would benefit from the established performance of STED microscopy in 3D cellular imaging.

In this work, and inspired by the single-molecule localization microscopy method DNA points accumulation for nanoscale topography (DNA-PAINT) ^15^, we employed DNA-fluorophore labels for intensity-based quantitative STED. DNA-fluorophore labels were previously used for STED microscopy and enabled bypassing photobleaching ^16^. The brightness could be enhanced by placing two fluorophores on a single DNA strand ^17,18^. Building on these concepts, and additionally employing locked nucleic acids (LNAs) ^19^, we designed DNA-fluorophore labels with four fluorophores that showed a stable intensity readout. We then used DNA origami as a molecular platform to demonstrate molecular counting of 1 to 7 targets reliably. In addition, we demonstrate quantitative STED in the cellular context by measuring the oligomeric state of the epidermal growth factor receptor (EGFR) in resting and EGF-treated cells. This approach paves the way for quantitative measurements that can be implemented on any accessible STED setup without complex technological demand.

## Results

First, we developed a robust strategy for protein labeling that would enable obtaining molecular counts from intensity measurements using STED microscopy. For this purpose, we designed a 44 nucleotide (nt) DNA backbone (‘docking strand’) to which four fluorophore-labeled sequence-complementary 11 nt DNA strands (‘imager strands’) can hybridize (**Figure 1A**). To increase the hybridization strength, we exchanged single nucleotides with their locked nucleotide counterpart (LNA) ^19^ (**Table S1**). The rationale for this label design was to (i) achieve a higher brightness per docking strand while using reasonable excitation intensity, and (ii) to minimize the impact of fluorescence intensity fluctuations that would occur for single fluorophores and impact the accuracy of the analysis.

**Figure 1:**
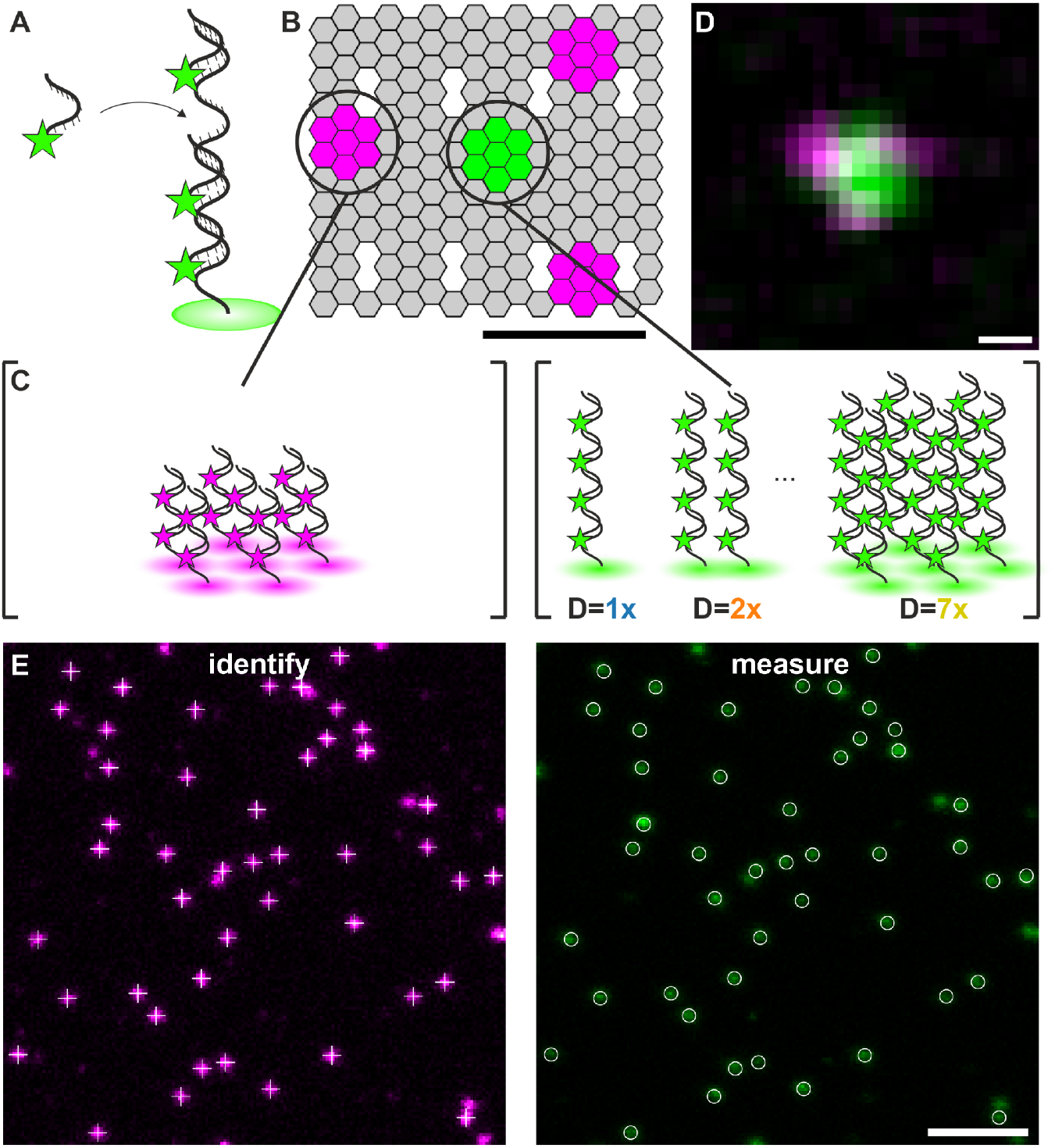
Design of a DNA origami platform and DNA-fluorophore labels for quantitative STED microscopy. **(A)** Design of a DNA/LNA hybrid (44 nt) carrying 4 fluorophores. A single-stranded DNA docking strand with 4x repeated oligonucleotide sequence binds up to four fluorophore-labeled DNA/LNA imager strands. (**B)** DNA origami design for two-color STED microscopy calibration experiments (Scale bar = 30 nm), carrying **(C)** 7 docking strands (22 nt) at three corner sites, with each docking strand binding two 11 nt imager strands labeled with Abberior STAR 635P (magenta, left). In the center, the target docking strand (44 nt) is positioned and binds four 11 nt DNA/LNA imager strands labeled with Abberior STAR 580 (green, center). The number of target docking strands varied between 1 and 7. (**D)** Exemplary two-color STED image of a single DNA origami (Scale bar = 50 nm). (**E)** DNA origami are identified using the fluorescence signal of Abberior STAR 635P as a reference (magenta, left image) and measuring the fluorescence intensity of the center peak (green, right image) (Scale bar = 1 µm).

We next tested the performance of the DNA/LNA-fluorophore probe for intensity-based molecular quantification by adapting a previously reported DNA origami platform ^20^ equipped with two different docking strand sequences (**Figure 1B**): the first docking strand (22 nt) was positioned at three corner sites (7 copies at each site), forming an asymmetric triangle that serves as a marker; the targeting docking strand (44 nt) was positioned at the center of the DNA origami, and varied between 1 and 7 copies (**Figure 1C**). Assuming a flat DNA origami structure and 5 nm spacing between staple strands, the center-to-corner distance was calculated to range from 27 to 31 nm, and the corner-to-corner distance from 46 to 56 nm (**Figure S1A**). The corner sites were targeted by 11 nt DNA imager strands labeled with Abberior STAR 635P, and the target docking strand at the center of the origami was targeted by 11 nt DNA/LNA imager strands labeled with Abberior STAR 580. Two-color STED images were recorded, and the structure of the DNA origami was resolved (**Figures 1D, S1B, C**). We localized the center of each position and measured the distances between the different sites on the DNA origami, yielding 36 ± 14 nm for the center-to-corner distance and 64 ± 8 nm for the corner-to-corner distance (**Figure S1D, S1E; Table S2)**. Using this DNA origami as a platform, we recorded 2-color STED images, used the triangular structure in the red spectral channel as an identifier, and read out the fluorescence intensity for a varying number of target docking strands in the center (**Figure 1E**). To assess the possible influence of chromatic aberration, we recorded STED images of gold beads in the two spectral channels and measured their pairwise distance to 4.0 ± 0.1 nm (**Figure S2**).

Next, we designed 6 different DNA origami with a varying number of target docking strands (1x, 2x, 3x, 4x, 5x, and 7x) and recorded two-color STED images (**Figure 2A**). An increase in the fluorescence signal with an increasing number of docking strands is clearly visible in the images. We performed a quantitative read-out of the fluorescence intensity of the target docking strand in the center of the DNA origami and generated intensity histograms (**Figure 2B**). Plotting the peak intensity of each measurement against the number of target docking strands showed a clear linear trend (**Figure 2C, Table S3**), with the y-axis intercept representing the background intensity. The width of the intensity distributions (obtained from Gaussian fitting) scales with the square root of the number of docking sites (**Figure 2D, Table S3)**.

**Figure 2:**
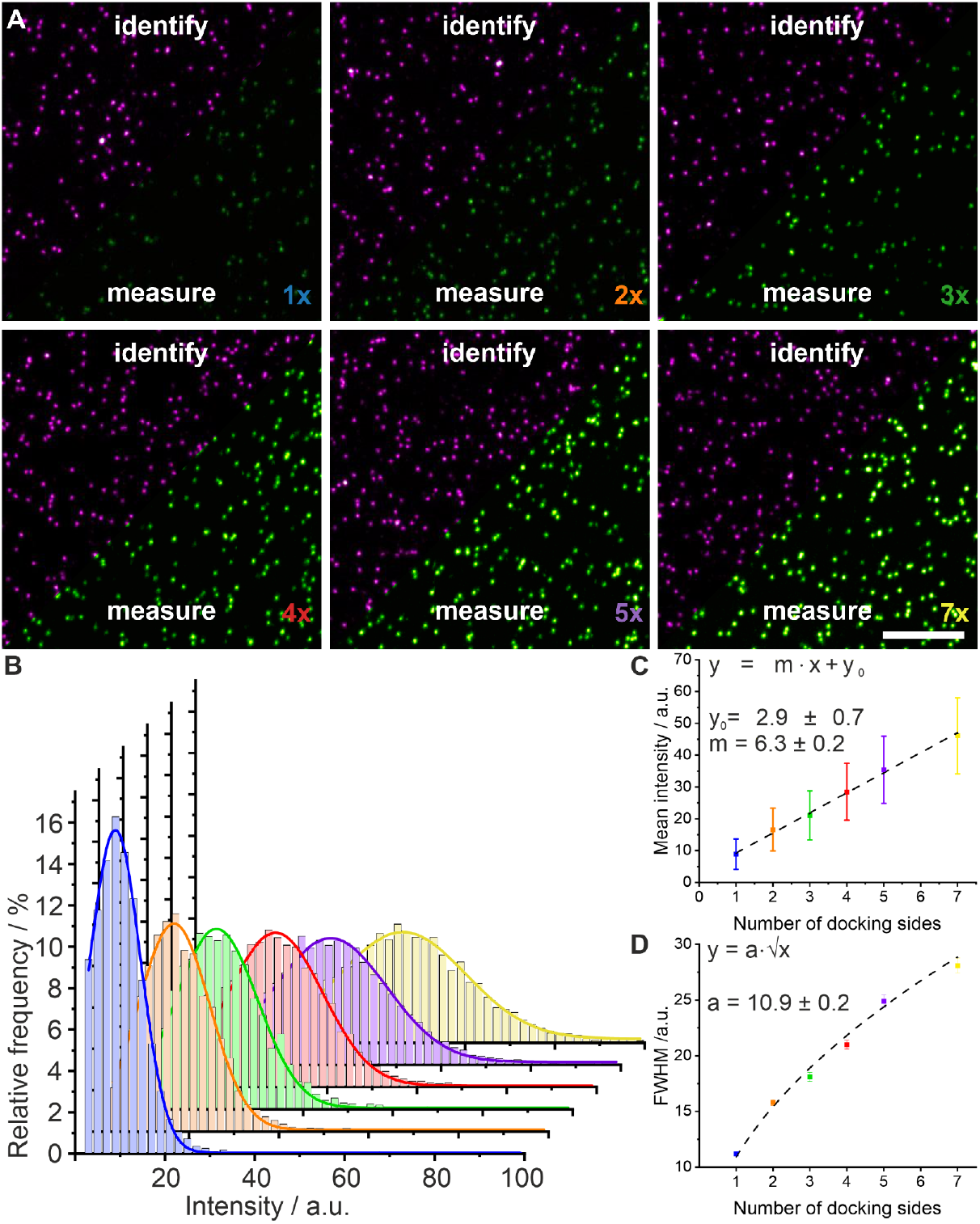
Quantitative STED microscopy of DNA Origami with a varying number of target docking strands. **(A)** Exemplary two-color STED measurements of six different DNA origami with a varying number of target docking strands, ranging from 1 to 7. The corner sites labeled with Abberior STAR 635P were used for identification (magenta), and the fluorescence intensity of the target docking strand in the center (green) was measured (scale bar 3 µm). (**B)** Intensity distributions for six different DNA origami carrying 1, 2, 3, 4, 5, or 7 target docking strands were fitted with a Gaussian function. **(C)** Peak intensities (squares) and FWHM (whiskers) from the histograms in (B) are plotted against the number of target docking strands. Linear regression (dotted line) was used to fit the data. **(D)** The FWHM is plotted against the number of target docking strands (errors of the fit are indicated as whiskers). A square root function (dotted line) was fitted to the data.

We next evaluated whether mixtures of the DNA origami containing different numbers of target docking strands can be discriminated within the same sample, to assess the resolving power of our approach for quantitative STED. For that purpose, we immobilized equal amounts of DNA origami containing two (1x and 2x, 1x and 3x, 1x and 4x, or 1x and 5x) or three (1x + 4x + 7x) target docking strands and performed two-color STED microscopy (**Figure 3A-F)**. Visual inspection of the STED images recorded for these mixtures clearly showed two or three different intensities for the central spot (magnified views in **Figure 3Bii-Fii**). For image analysis, we fitted the intensity distributions with either two (**Figure 3Biii-3Eiii**) or three Gaussians (**Figure 3Fiii**) using FWHM values from the calibration data (**Methods, Table S3**). The results of this analysis clearly retrieved the fraction and number of target docking strands in the respective sample (**Table S4**). We note that although the intensity distribution of 1x + 2x was not bimodal, we were still able to identify the correct number of target docking strands and fractions (**Table S5**). In summary, the peak centers determined from mixtures of DNA origami with different numbers of target docking strands corresponded well to the values obtained for single DNA origami (**Table S3, Table S4**). Therefore, our approach is suitable for extracting molecular numbers from super-resolved clusters using intensity information from STED microscopy images.

**Figure 3:**
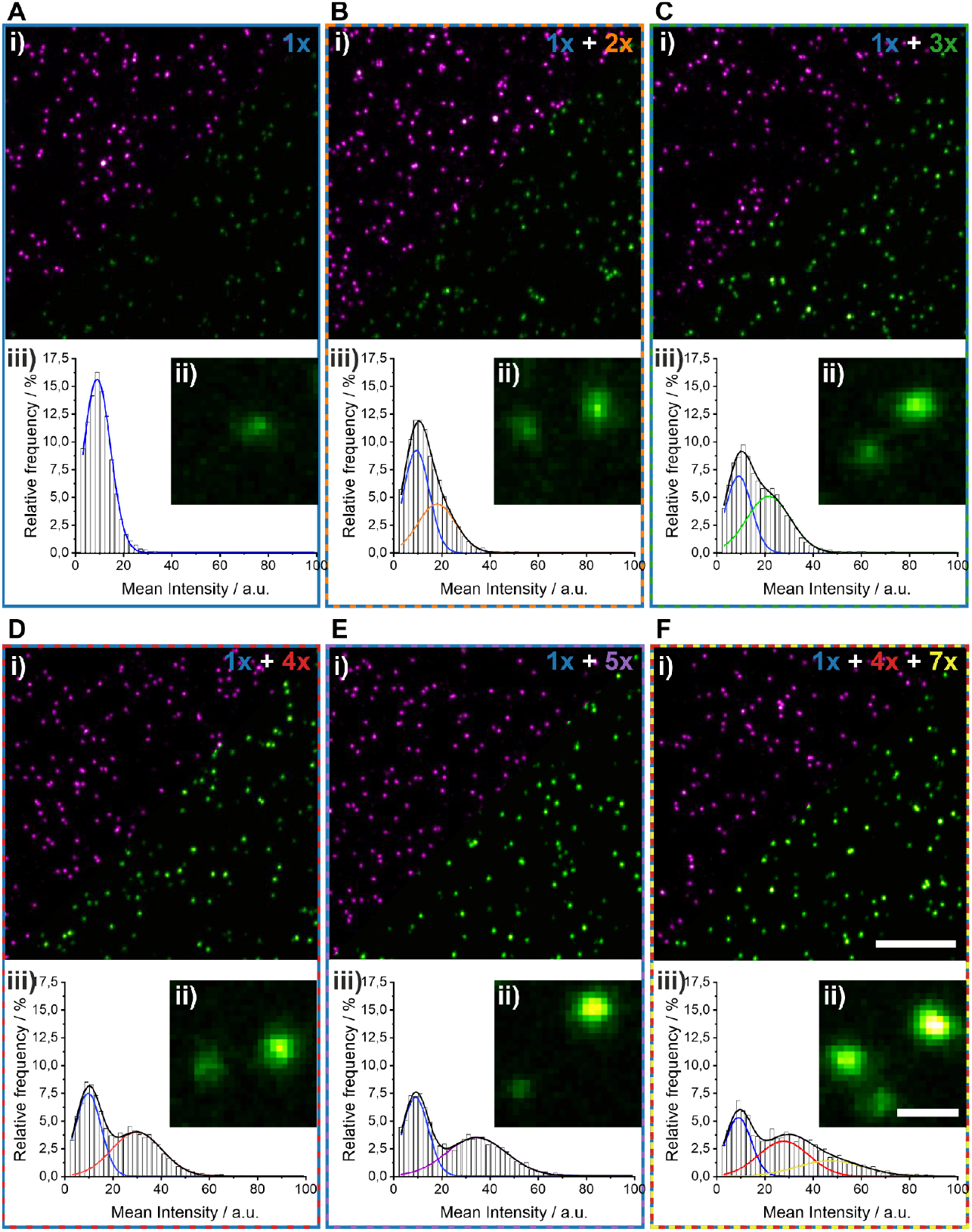
Quantitative STED microscopy of mixtures of DNA origami with different numbers of target docking strands. **(A)** Two-color STED microscopy of a DNA origami containing one target docking strand, **(B-E)** a mixture of two origami with a different number of target docking strands, and **(F)** a mixture of three DNA origami with 1, 4, and 7 target docking strands. In each panel, (i) an overview image (Abberior STAR 635P, magenta; Abberior STAR 580, green) is shown (scale bar 3 µm), (ii) a magnified region (scale bar 300 nm), and (iii) intensity distributions with Gaussian fit functions.

Next, we evaluated whether this approach can be used to quantify protein numbers of super-resolved nanoclusters in fixed cells. For this purpose, we chose the epidermal growth factor receptor (EGFR), a membrane receptor that is reported to be mainly monomeric in resting cells and dimerizes upon ligand binding ^21–23^ (**Figure 4A**). Transiently transfected CHO cells expressing EGFR-ALFA-mEGFP were labeled with two nanobodies: a first one, labeled with Abberior STAR 635P and targeting mEGFP (NB@GFP-STAR 635P); a second one, carrying a single 44 nt target docking strand (NB@ALFA-4xP3) and targeting the ALFA-tag ^24^ (**Figure 4A, left**). Two-color STED microscopy was performed in resting and EGF-treated cells (**Figure 4B**). For quantitative analysis, the fluorescence signal of Abberior STAR 635 (targeting GFP) served as a reference signal, and the fluorescence signal of Abberior STAR 580 bound to the target docking strand (targeting the ALFA-tag) was used for quantitative analysis. This two-step approach reduces the contribution of unspecific signals to the analysis, which might occur if only a single label was used. The resulting intensity distribution for resting cells (**Figure 4Ci**) shows a bimodal distribution: the first peak at 2.0 ± 0.1 a.u. corresponds to the background signal, when no ALFA-tag targeting nanobody is bound; the second peak at 7.0 ± 0.8 a.u. corresponds to the ‘true signal’ for monomeric EGFR (**Table S6**), and is similar to the data recorded for a single target docking strand on DNA origami (**Figure 2C, D, Table S3**). Furthermore, this enabled us to determine the labeling efficiency of NB@ALFA-4xP3 (see Methods) ^25^ to 59 ± 9 % (**Figure S3**). In EGF-treated cells (see Methods), the intensity distribution showed a tailing to higher intensities (**Figure 4Cii**), indicating oligomerization of EGFR. Assuming only monomers and dimers for EGFR, we fitted the intensity histogram with 3 Gaussian functions (see Methods) and found intensity values of 2.7 ± 0.1 a.u. (background), 6.4 ± 0.9 a.u. (monomer and single-labeled dimer) and 16 ± 3 a.u. (dual-labeled dimer) (see Methods) (**Figure S4A, Table S6**). Additionally, we observe a labeling efficiency corrected EGFR dimer fraction of 55 ± 7 % (**Figure 4Ciii, Figure S4B, Table S7**).

**Figure 4:**
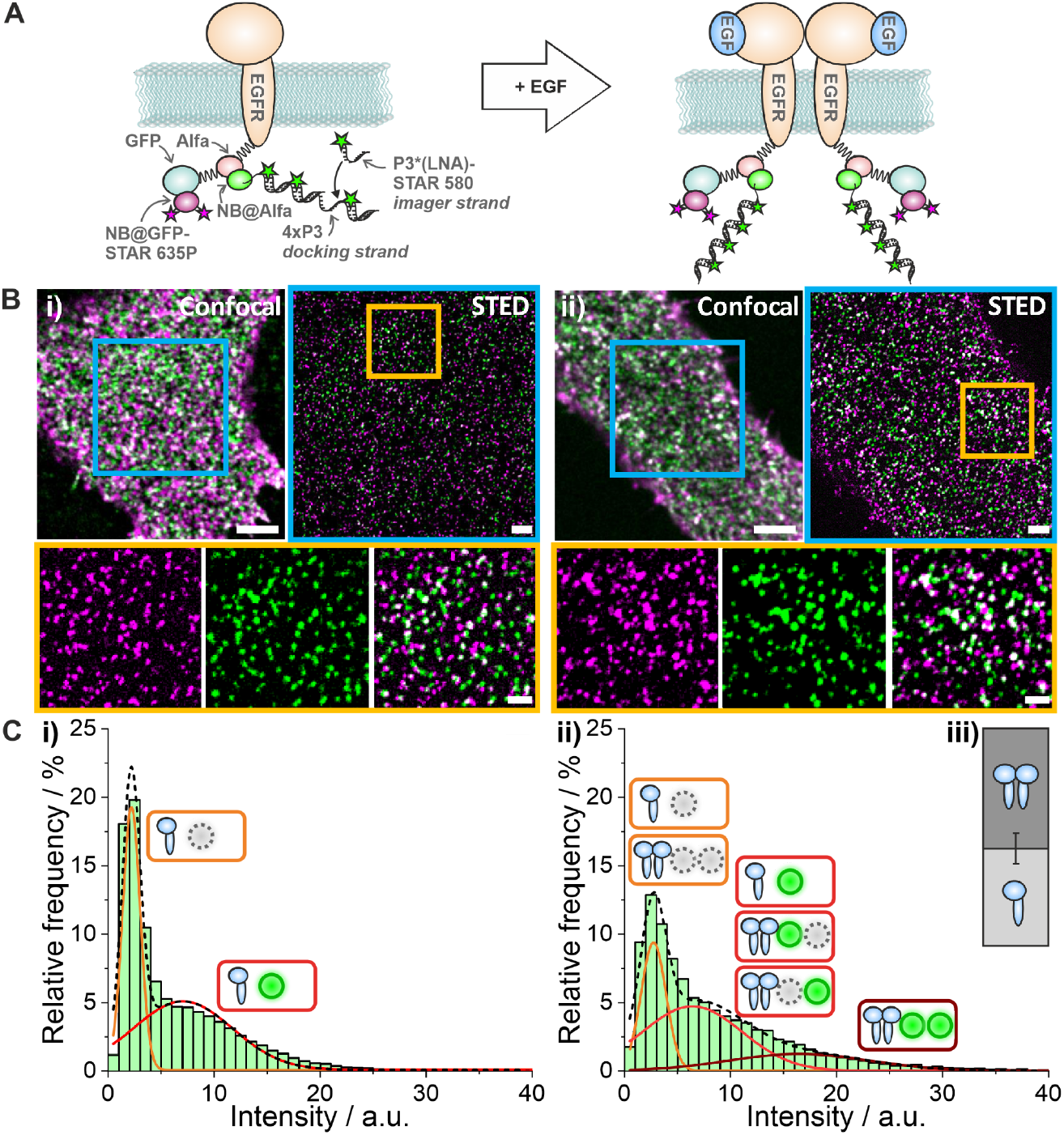
Quantitative STED microscopy of EGFR dimerization in CHO cells. **(A)** Schematic illustration of EGFR-ALFA-mEGFP fusion protein (left) and canonical model of EGFR dimerization (right). **(B)** Two-color confocal and STED microscopy of EGFR labeled with NB@GFP-STAR 635P (magenta) and NB@ALFA-4xP3 (green) in a resting cell **(i)** and an EGF-treated cell **(ii)** with magnified views (below). Scale bars are 4 µm (confocal overview), 2 µm (STED overview), 500 nm (magnified STED images). **(C)** Intensity distributions for **(i)** resting and **(ii)** EGF-stimulated cells, with Gaussian fits and small icons showing contributions to the individual peaks (see Methods). **(iii)** Corrected EGFR dimer fraction in EGF-stimulated cells (see Methods) and standard error indicated as a whisker.

We quantified the cluster density of EGFR per area using the NB@GFP-STAR 635P signal to 5.7 ± 1.3 µm^-2^ (resting) and 5.4 ± 1.5 µm^-2^ (EGF-treated), respectively (**Table S8**). Additionally, negative controls were performed in untransfected cells, yielding a cluster density of 0.2 ± 0.1 µm^-2^ (NB@GFP-STAR 635P) and 0.4 ± 0.3 µm^-2^ (NB@ALFA-4xP3) (**Figure S5A, Table S8**). Furthermore, we found that pre-incubation of NB@ALFA-4xP3 with the DNA/LNA-Abberior STAR 580 imager strands strongly decreased the degree of unspecific binding to cells (**Figure S5B, Table S8**). Since STED is a microscopy technique that is ideal for recording large-volume 3D super-resolution data of cell biology samples ^4^, we also performed volumetric STED imaging of EGFR in an entire CHO cell in 3D (**Figure S6**).

## Discussion

We report quantitative STED microscopy and demonstrate molecule counting with single-protein resolution and in super-resolved clusters. Inspired by DNA-based fluorophore labels in SMLM ^15,17^, we first designed a DNA/LNA-based label that carries 4 fluorophores in order to achieve signal amplification. This is a key prerequisite for intensity-based quantitative analysis, bypassing the uncertainties of single-fluorophore intensity analysis or the need for sophisticated instrumentation ^14^ and enabling single-target detection. We demonstrated robust intensity-based analysis of 1 to 7 fluorophore labels (**Figure 2**), of mixtures (**Figure 3**), and of the oligomeric state of EGFR receptors in cells (**Figure 4**). The probe design and intensity-based analysis are easy to implement on any basic STED microscope without the need for sophisticated technical additions, representing a valuable extension to this powerful microscopy method.

One of the major applications of quantitative super-resolution microscopy is the extraction of the oligomeric state of nanoscale protein clusters in cells. The assembly of proteins to nanoscale clusters is linked to a shift in the functional state, for example, in the dimerization of receptor tyrosine kinases ^26,27^ and G-protein coupled receptors ^28^, the oligomerization of tumor necrosis factor receptors ^29^, and clustering of T-cell receptors ^30^. So far, microscopic studies addressing nanoscale protein clustering mostly used single-molecule super-resolution methods ^9^, in particular approaches that relate single-molecule emission kinetics to molecule numbers ^10,11^. However, this requires sophisticated data analysis, it contains uncertainties since single-molecule emission kinetics are affected by the local environment, and it is limited by the imaging characteristics of SMLM methods, such as long acquisition times and limited 3D range. Other high-end microscopy techniques, such as MINFLUX ^13^ and resolution enhancement by sequential imaging (RESI) ^31^, enable the direct visualization of proteins within a dense nanoscale assembly. However, this impressive spatial resolution comes at the cost of complex instrumentation, very long acquisition times, and restricted observation volumes.

Here, we demonstrated fast quantitative STED and measured the oligomeric state of EGFR in the plasma membrane of resting and EGF-treated CHO cells with single-protein resolution. The molecular mechanism of EGFR activation is well understood: upon ligand binding, EGFR dimerizes, thereby initiating transphosphorylation, which in turn recruits and activates key downstream signaling effectors ^32,33^. Previous work showed that EGFR predominantly is monomeric in resting cells, and assembles into dimers and higher-order oligomers upon EGF stimulation ^27,34^. Using quantitative STED, we reliably measured fractions of monomeric and dimeric EGFR that agree well with these previous results. In order to assess the precision of our approach, we employed a published EGFR expression system ^27^ enables dual-color labeling of EGFR through two specific tags at the C-terminus, mEGFP and ALFA-tag. This allowed us to quantify the labeling efficiency of the anti-ALFA nanobody, which is needed for absolute quantification. We modified a reported procedure to determine the labelling efficiency by additionally using the intensity information (**Figure 4Ci**) and hereby decoupling it from manual thresholding ^25^. With that, we found a labeling efficiency of 59% for the anti-ALFA nanobody that is higher than previously reported ^27^, which might be related to the shorter illumination time as compared to DNA-PAINT, and thus less photo-damage of docking strands ^35^. In addition, we found a much reduced unspecific binding of the anti-ALF nanobody when pre-incubated with the DNA/LNA-fluorophore labels before the addition to cell samples. Using the dual-labeling approach had thus two major benefits: first, our analysis focused on true signal and minimized the impact of unspecific signal; second, we were able to directly correct the EGFR dimerization ratio for the labeling efficiency, which would not be possible with a single-color approach.

In summary, we present a robust approach for intensity-based molecular quantification using STED microscopy. The major advantages are that this approach is fast, easy to implement and analyze, does not require complex instrumentation, and profits from the imaging opportunities provided by STED microscopy such as 3D, large volume, and multi-color imaging. The probe design can be extended and adapted in brightness, color, and for multiple targets, e.g. by integrating DNA barcoding. Given the overall simplicity of the approach, we envision a low entry-barrier for integration of this method into a broad spectrum of microscopy experiments in cell biology.

## Methods

### Sample preparation

#### Folding and purification of DNA origami

A rectangular DNA origami structure was designed using the Picasso design tool. ^15^. Annealing of the DNA strands was performed in a mixture of 40 µL 1x TE (Tris-EDTA) buffer with 12.5 mM MgCl2 containing 10 nM scaffold strand (M13mp18; tilibit nanosystem, Germany), 100 nM core staple strands, 1 μM biotinylated staple strands, and 1 μM docking strand handles. The mixture was incubated for 5 min at 65°C and subsequently cooled to 25°C over the course of 4 h. The DNA origami was purified using 100 kDa MWCO ultra centrifugal filter units (Amicon Ultra, Germany) and stored in FoB5 buffer (5 mM Tris pH 8, EDTA pH 8, 5 mM NaCl, 5 mM MgCl_2_) at -20°C. For strand sequences, see **Tables S9 - S11**.

#### Immobilization of DNA origami

The channel of a channeled glass slide (µ-Slide VI 0.5 Glass Bottom, Ibidi, Germany) was filled with 1x PBS. Subsequently, biotinylated BSA (1 mg/mL, dissolved in 1x PBS) was added and incubated for 15 min. After 3x washing with 1x PBS, streptavidin (0.2 mg/mL, dissolved in 1x PBS) was added and incubated for 15 min. The channel was washed 3x with 1x PBS, and the DNA origami solution was added. After incubation for 20 min, the channel was washed 3x with an imaging buffer (1x PBS, 500 mM NaCl). For imaging, 20 nM P2 and P3 imager strands (**Table S1**) were diluted in PBS with 500 mM NaCl and added right before the experiment.

#### Cell culture, EGFR transfection, and stimulation

Chinese hamster ovary cell line (CHO-K1) wild type (WT) (Lonza) was grown in growth medium consisting of high glucose Dulbecco’s modified Eagle medium/nutrient mixture F-12 (DMEM/F12, Gibco, Life Technologies, USA), supplemented with 1% GlutaMAX (Gibco, Life Technologies) and 10% fetal bovine serum (FBS) (Gibco, Life Technologies) under humidified conditions at 37°C and 5% CO_2_. All cells were passaged every 3-4 days or upon reaching 80% confluency. 10000 cells per well were seeded in 8-well chambers (SARSTEDT AG & Co. KG, Nümbrecht, Germany) coated with PLL-PEG-RGD (prepared according to Li et al. ^26^) 2 days before transfection. Directly before transfection, the cells were washed with growth medium once and transfected using jetOPTIMUS® DNA Transfection Reagents (SARTORIUS, Germany). The transfection process was carried out in accordance with the protocol established by the supplier without additional explanation. In brief, 0.5 µg of the plasmid containing the EGFR-ALFA-EGFP construct (kind gift from Prof. Ralf Jungmann) ^27^ was diluted in 50 µL jetOPTIMUS buffer and mixed with 0.5 µL reagent for each well transfection. The plasmid mixture was replaced by DMEM/F12 medium 4 h after addition. The cells were allowed to express EGFR for 20 h. For stimulation, cells were treated with 100 ng/mL epidermal growth factor (EGF, #E9644, Sigma-Aldrich, Germany) at 37°C for 1 min, subsequently washed with 0.4 M sucrose (Sigma-Aldrich, Germany) in 1x PBS, and fixed with 4% FA in 1x PBS at 37 °C for 15 min.

#### Immunofluorescence labeling

For immunolabeling, CHO cells were incubated using an antibody-incubation buffer (Massive Photonics, Germany) for 30 min at room temperature. Cells were washed three times with PBS and incubated with an antibody incubation buffer containing FluoTag-x4 anti-GFP Abberior STAR RED nanobodies (NanoTag Biotechnologies, Germany) and alpaca anti-ALFA nanobodies modified with the custom-designed 4xP3 sequence (Massive Photonics, Germany) (**Table S1**) in a 1:100 dilution for 1 h at room temperature. After washing three times with PBS, cells were post-fixated using 4% FA in PBS for 10 min. For imaging, 20 nM P3-LNA (**Table S1**) was diluted in PBS with 500 mM NaCl and added right before the experiment.

### STED Microscopy

All imaging experiments were conducted with an Abberior expert line microscope (Abberior Instruments, Germany), operated either in confocal or STED imaging mode. The microscope was equipped with an Olympus IX83 body (Olympus Deutschland GmbH, Germany) and a UPLXAPO 60x NA 1.42 oil immersion objective (Olympus Deutschland GmbH, Germany).

#### Imaging of DNA origamis

For STED image acquisition of DNA Origami, samples were excited with either a 561 nm or 640 nm pulsed excitation laser (3.7 µW and 3.2 µW at the back focal plane) and depleted using a 775 nm pulsed laser (157 mW and 109 mW at the back focal plane) having a 2D doughnut point spread function. All lasers had a pulse repetition rate of 40 MHz, and the fluorescence was detected 750 ps after every pulse over a width of 8 ns. Fluorescence was collected in the spectral range of 571-630 nm (561 nm excitation) and 650-763 nm (640 nm excitation) using two APDs. The images were acquired with a pinhole of 1.0 AU, line accumulation of 25 (561 nm excitation) and 20 (640 nm excitation), pixel dwell time of 5 µs, and a pixel size of 30 nm.

#### Imaging of CHO cells

Confocal imaging of CHO cells was performed using a 561 nm excitation laser (3.3 µW at the back focal plane) and a 640 nm excitation laser (3.2 µW at the back focal plane). Fluorescence was collected in the spectral range of 571 nm to 630 nm, respectively, 650 nm to 763 nm using two avalanche photodiodes (APD). The images were acquired with a pinhole of 1.0 AU, a pixel dwell time of 5 µs, and a line accumulation of 10 (561 nm excitation) and 5 (640 nm excitation). A pixel size of 50 nm was used.

For STED imaging of CHO cells, samples were excited with either a 561 nm or 640 nm pulsed excitation laser (3.3 µW and 3.2 µW at the back focal plane) and depleted using a 775 nm pulsed laser (109 mW and 91 mW at the back focal plane) having a 2D doughnut point spread function and with an excitation delay of 750 ps and a 8 ns width. Fluorescence was collected in the spectral range of 571-630 nm (561 nm excitation) and 650-763 nm (640 nm excitation) using two APDs. The images were acquired with a pinhole of 0.81 AU, line accumulation of 35 (561 nm excitation) and 20 (640 nm excitation), respectively, pixel dwell time of 5 µs, and a pixel size of 30 nm.

For volumetric STED imaging of CHO cells, samples were excited with either a 561 nm or 640 nm pulsed excitation laser (1.5 µW and 3.2 µW at the back focal plane) and depleted using a 775 nm pulsed laser (45 mW and 28 mW at the back focal plane) having a 3D top hat point spread function and with an excitation delay of 750 ps and a 8 ns width. Fluorescence was collected in the spectral range of 571-630 nm (561 nm excitation) and 650-763 nm (640 nm excitation) using two APDs. The images were acquired with a pinhole of 0.71 AU, line accumulation of 7, pixel dwell time of 17 µs, and a pixel size of 60 nm.

Imaging settings for each figure are summarized in **Table S12**.

### Image analysis

#### Intensity-based molecular counting

Molecular counting was performed by determining the intensity of the 4x-labeled DNA docking strand, either at the center of DNA origami (**Figures 1, 2, 3**) or attached to a nanobody (**Figure 4**). Images were analyzed using ImageJ ^36^. The signal maxima in the fluorescence channel of STAR 635P (**Figure 1B, C; Figure 4A, left**) were identified using the „Find maxima” plugin. For this, a threshold (“prominence”) of 15 was used for DNA origami, and 10 for EGFR receptors. Pixel intensities in the STAR 580 channel were measured at the identified coordinates. Frequency of intensity values was counted using a bin size of 2 a.u. for DNA origami and 1 a.u. for EGFR.

In the case of DNA origami, a threshold of 2 was used to exclusively analyse the target-specific signal. The percentage of the signal analyzed from the total signal is given in **Table S13**. Intensity histograms were fitted using a Gaussian distribution

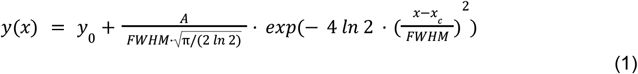

with offset *y*_*0*_, center *x*_*c*_, area of the integral A, and the width at half height *FWHM*. In experiments with 2 or 3 different DNA origami containing different numbers of docking strands (**Figure 3**), a respective number of Gaussian functions were used to fit the intensity distributions, and FWHM values were fixed to the numbers obtained from the respective experiment with unmixed DNA origami (**Table S3**). Fractions of Gaussian distributions *f*_*i*_ were calculated as

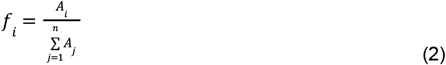

With *A*_*i*_ being the corresponding area of the Gaussian distribution integral and 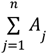 the sum of all areas. The data was anonymized, and the user fitting the histogram data did not have knowledge of the composition of the origami used. Recovered fraction ratios for the DNA origami mixtures are given in **Table S4**.

In multi-origami samples where a clear separation was difficult to recognize (**Figure 3**), e.g., in mixtures of DNA origami containing one and two docking strands (1x + 2x) or 3 different docking strands (1x + 4x + 7x), the accuracy of fitting the intensity distribution was evaluated by varying center and FWHM values (**Table S5**).

Intensity histograms obtained for EGFR in CHO cells were fitted using Gaussian distributions (equation (1); **Table S6**). For resting cells, 2 Gaussian distributions were used. Fractions *f*_*i*_ were calculated using equation (2). To determine the labeling efficiency of the ALFA-tag targeting nanobody NB@ALFA-4xP3, we followed a published procedure ^25^ and assumed that EGFR predominantly occurs as monomers in resting cells based on literature ^27^. We fitted the intensity distribution (**Figure 4Ci**) with two Gaussian functions to determine the fractions of background signal (1st peak) and DNA/LNA-fluorophore-labeled NB@ALFA-4xP3 (2nd peak; **Figure S4**). From this, we could calculate the labeling efficiency *LE:*

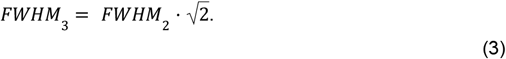

Using these values, the labeling efficiency was calculated to be 59 ± 9 %.

For stimulated cells, 3 copies of the functions were used. For this, the *FWHM* of the second peak was fixed to the value obtained for the second peak in resting cells, and the *FWHM* value for the third peak was calculated using

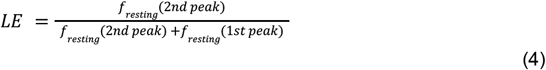

The results of the 3 Gaussian fitting include peak centers, FWHMs, and the fractions are reported in **Table S6**. For stimulated cells with mostly monomeric and dimeric EGFR ^27^, the measured fractions f can be expressed as the following three equations:

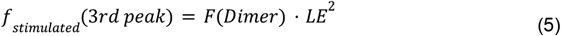

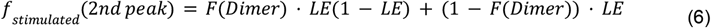

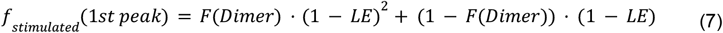

only dependent on the corrected Dimer fraction *F(Dimer)* and the labeling efficiency *LE*.

Using these, the corrected dimer fractions were calculated as 56 ± 8 % (via first peak fraction), 52 ± 5 % (via second peak fraction), and 58 ± 17 % (via third peak fraction) and a mean value of 55 ± 7 % (**Table S7**).

#### Cluster density

To extract receptor density, images were analyzed using ImageJ ^36^. At first, a mask encompassing the cell shape was drawn. Receptors were counted within the mask using the plugin “Find maxima” using a threshold of 10 a.u. for STAR 635P and 7 a.u. for STAR 580. The density was calculated as the number of receptors divided by the area of the corresponding mask (Results in **Table S8**)

## Data availability

All imaging data are available in an online repository, DOI: https://doi.org/10.5281/zenodo.17522871

## Author contributions

L.F.K. and M.H. designed the project. L.F.K. performed microscopy experiments, Y.L. prepared cell samples. All authors analyzed and discussed the data and wrote the manuscript.

## Acknowledgement

We thank Petra Freund for assistance with cell culture. We acknowledge funding by the Deutsche Forschungsgemeinschaft (CRC1507, project-id: 450648163 (M.H., Y.L.); CRC 1177, project-id: 259130777 (M.H., A.B.); INST 161/1020-1 FUGG (M.H.)); the SubCellular Architecture of LifE (SCALE) consortium funded by the Goethe University Frankfurt, Germany (M.H., L.F.K.); the GRADE Center SCALE-it (L.F.K., Y.L.).

