## Supplementary Information for "Quantitative STED microscopy with DNA-fluorophore labels"

Laurell F. Kessler[1], Yunqing Li[1], Ashwin Balakrishnan[1], Mike Heilemann[1,#]
[1]Institute of Physical and Theoretical Chemistry, Goethe-University Frankfurt, Max-von-Laue-Strasse 7, 60438 Frankfurt, Germany

### SUPPLEMENTAL INFORMATION

**Supplemental Figures S1 - S6**
**Supplemental Tables S1 - S13**

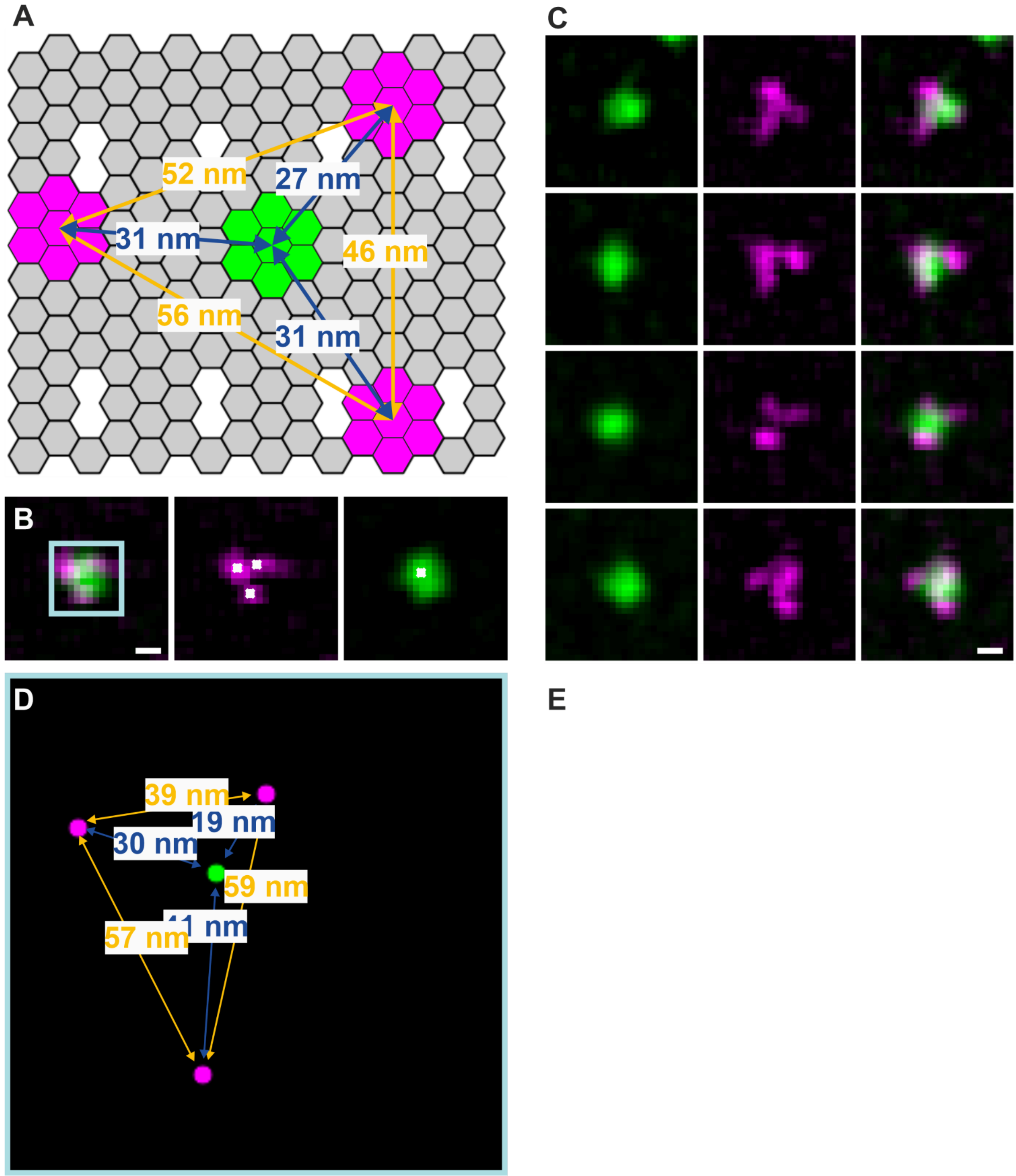


**Figure S1: STED imaging of DNA origami and distance measurement. (A)** Folding scheme of DNA origami and calculated distances assuming a hexagon size of 5 nm. (**B)** Exemplary two-color STED measurement of a single DNA origami (composite image, left; outside triangle, middle; center, right) with localized maxima indicated as white crosses. Scale bar = 50 nm. (**C)** Further examples of STED images of single DNA origami. Scale bar = 50 nm. (**D**) Zoom-in of the boxed region in (B) and inter-site distances. (**E)** Manual measured center-to-edge (blue) and edge-to-edge (yellow) distances from 5 DNA origami (B, C). Black lines indicate the median values, the squares indicate the mean values, the box indicates the 25th and 75th percentiles, and the whiskers indicate the minimum and maximum.

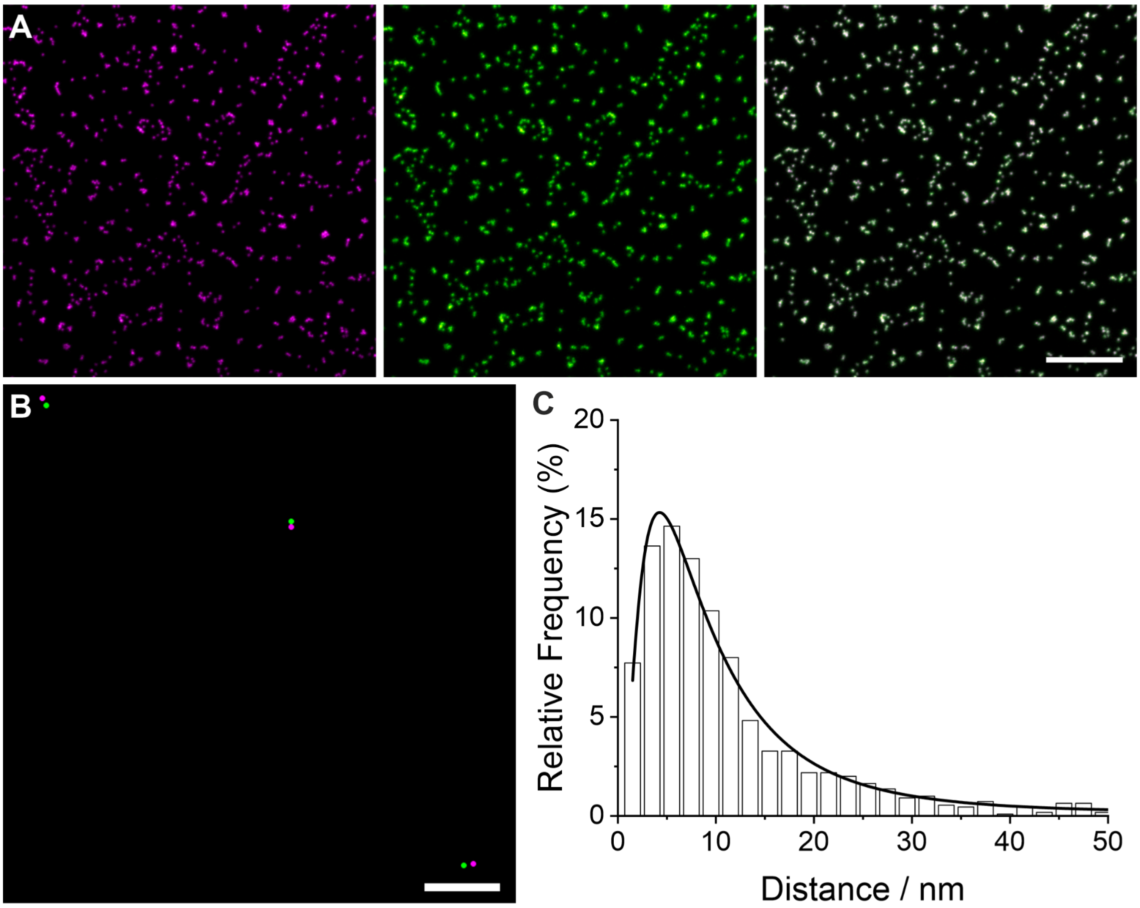


**Figure S2: Chromatic aberration of the STED microscope assessed by measuring gold beads. (A)** Two-color STED microscopy of gold beads on a glass surface using a 640 nm excitation laser (magenta; left) and a 561 nm excitation laser (green, middle). A composite of both channels is shown on the right. Scale bar = 3 µm. (**B)** Magnified view showing localized maxima of both channels. Scale bar = 50 nm. **C** Distribution of the nearest neighbor distance. A log-normal function was used to fit the histogram, with a peak at 4.0 +/- 0.1 nm.

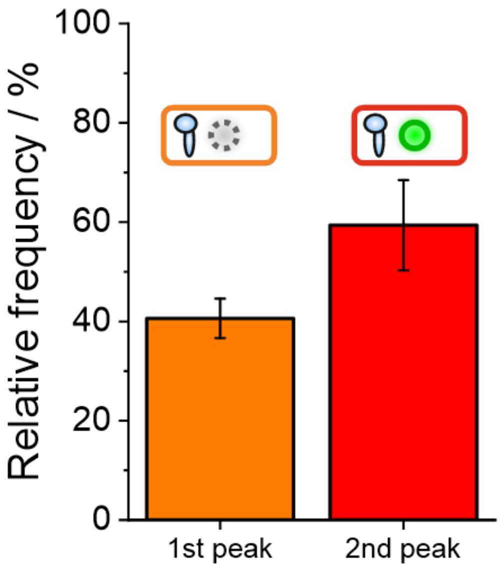


**Figure S3: Detection efficiency of the ALFA-tag targeting nanobody.** The 1st peak corresponds to the background signal, the 2nd peak corresponds to the fluorescence signal detected for NB@ALFA-4xP3. Relative occurrence was obtained by fitting two Gaussian distributions to the intensity distribution from resting cells (**Figure 4Ci**). From this, the correction factor for labeling was determined (equation (4), Methods).

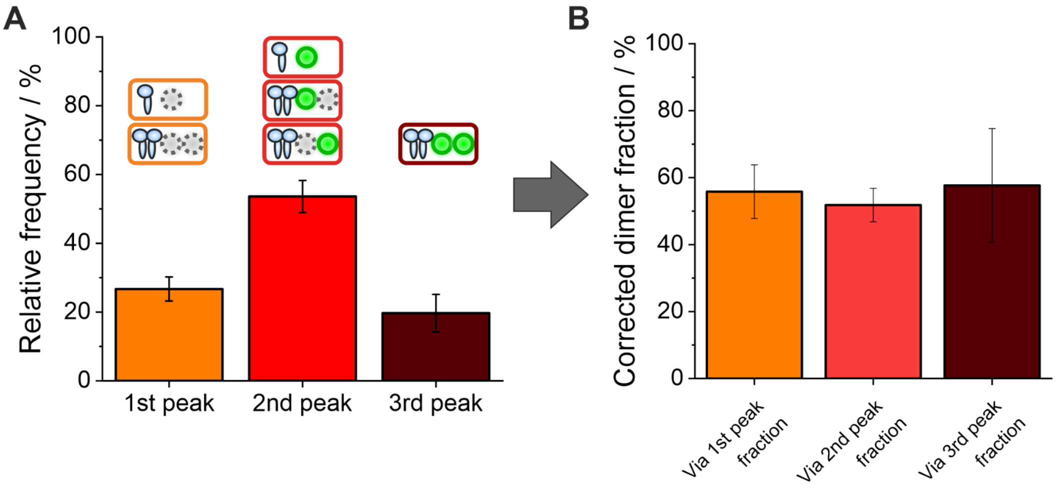


**Figure S4: Absolute quantification of EGFR dimerization. (A)** Normalized integrated intensity obtained by fitting 3 Gaussian distributions to the intensity histogram of EGF-stimulated cells (**Figure 4Cii**). The 1st peak corresponds to the background signal, the 2nd and 3rd peak are two intensity levels detected for NB@ALFA-P3. **(B)** Fraction of EGFR dimerization corrected for NB@ALFA-P3 labeling efficiency from 1st (equation (5)), 2nd (equation (6)), and 3rd peak (equation (7)) (Methods).

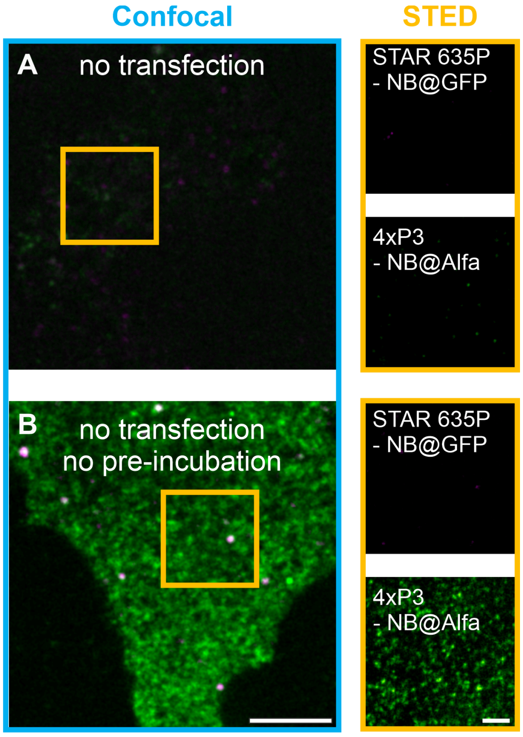


**Figure S5: Negative controls using non-transfected CHO cells. (A)** Exemplary untransfected cell measured with confocal and STED microscopy stained with NB@GFP-STAR 635P (magenta) and NB@ALFA-4xP3 (pre-incubated with P3-STAR 580; green). **(B)** Exemplary untransfected cell measured with confocal and STED microscopy stained with NB@GFP-STAR 635P (magenta) and NB@ALFA-4xP3 (no pre-incubation; green). Scale bar = 5 µm (confocal); 1 µm (STED).

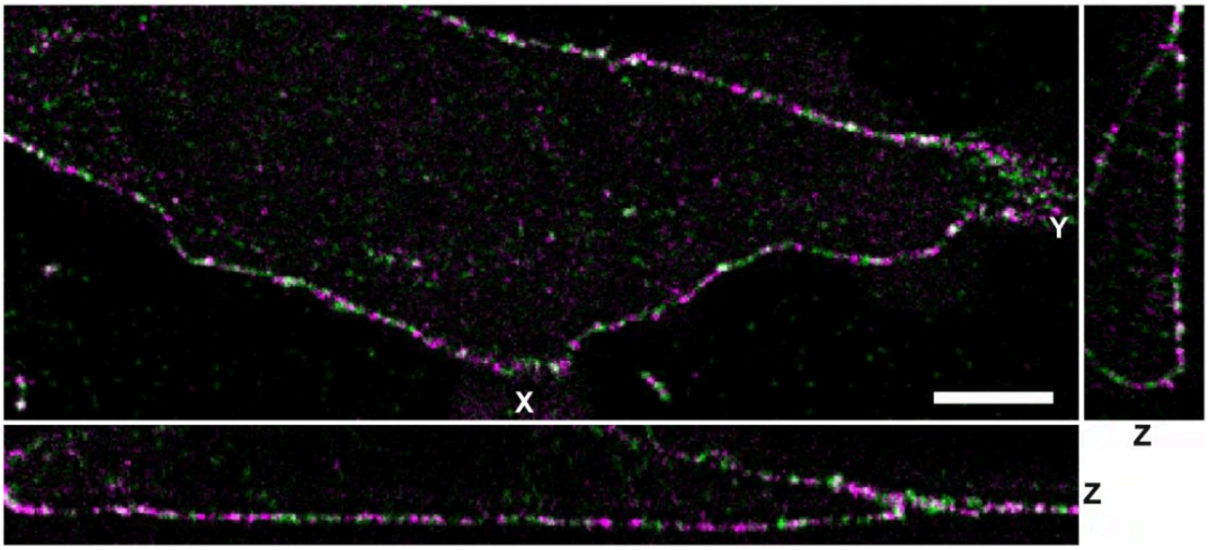


**Figure S6: Volumetric STED microscopy of EGFR in a resting CHO cell.** CHO cells expressing EGFR-ALFA-mEGFP were labeled with NB@ALFA-4xP3 (green) and NB@GFP-STAR 635P (magenta) (Scale bar = 4 µm).

### Supplemental Tables

**Table S1**: Imager and docking strand sequences used in this study. Modified LNA nucleotides are colored blue (Guo et al. 2019). Docking sequences were synthesized together with the origami folding sequences from 5' to 3' end. Conjugation of the 4xP3 docking strand to the nanobody against GFP was performed by Massive Photonics (Germany).

| **Name** | **Sequence** | **Supplier** |
|---|---|---|
| P2-STAR 635P | 5' TATG**TAG**ATAA-STAR 635P-3' | biomers.net |
| P3-STAR 580 | 5' TAA**TGA**AGAAA -STAR 580-3' | biomers.net |
| 2xP2 docking strand (Origami) | 5' *Origami folding sequence* - TTATCTACATATTATCTACATA - 3' | Eurofins Genomics |
| 4xP3 docking strand (Origami) | 5' *Origami folding sequence* - TTTCTTCATTATTTCTTCATTATTTCTTCATTATTTCTTCATTA - 3' | Eurofins Genomics |
| 4xP3 docking strand (Nanobody@GFP) | Nanobody - 5' - TTTCTTCATTATTTCTTCATTATTTCTTCATTATTTCTTCATTA - 3' | Massive Photonics |

**Table S2**: Calculated distances from two-color STED measurements of DNA origamis.

| **Pair** | **Distance / nm** |
|---|---|
| STAR 635P to STAR 580 (corner-to-center distance, magenta to green) | 36 ± 14 |
| STAR 635P to STAR 635P (corner-to-corner distance, magenta to magenta) | 64 ± 8 |

**Table S3**: Intensity analysis of DNA origami with different numbers of target docking strands. Errors are given as standard errors of the Gaussian fit.

| | **1x** | **2x** | **3x** | **4x** | **5x** | **7x** |
|---|---|---|---|---|---|---|
| Mean / a. u. | 8.9 ± 0.1 | 16.6 ± 0.1 | 21.1 ± 0.2 | 28.5 ± 0.2 | 35.4 ± 0.3 | 46.1 ± 0.2 |
| FWHM / a. u. | 11.2 ± 0.2 | 15.8 ± 0.3 | 18.1 ± 0.4 | 21.0 ± 0.4 | 24.9 ± 0.6 | 28.1 ± 0.7 |

**Table S4**: Fitting results for mixed DNA origami samples. Fixed values are indicated in red. Errors are given as standard errors of the Gaussian fit.

| | **1x+2x** | | **1x+3x** | | **1x+4x** | | **1x+5x** | | **1x+4x+7x** | | |
|---|---|---|---|---|---|---|---|---|---|---|---|
| | Peak 1 | Peak 2 | Peak 1 | Peak 2 | Peak 1 | Peak 2 | Peak 1 | Peak 2 | Peak 1 | Peak 2 | Peak 3 |
| Mean / a. u. | 8.9 | 16.6 | 9.1 ± 0.2 | 21.9 ± 0.5 | 9.6 ± 0.1 | 30.0 ± 0.4 | 9.0 ± 0.2 | 37.2 ± 0.5 | 8.6 ± 0.3 | 27.3 ± 1.1 | 45.9 ± 3.6 |
| FWHM / a. u. | 11.2 | 15.8 | 11.2 | 18.1 | 11.2 | 21.0 | 11.2 | 24.9 | 11.2 | 21.0 | 28.1 |
| Fraction / % | 54 ± 1 | 46 ± 1 | 47 ± 2 | 53 ± 2 | 50 ± 1 | 50 ± 1 | 49 ± 2 | 51 ± 2 | 34 ± 1 | 44 ± 4 | 22 ± 4 |

**Table S5**: Alternative fitting models for the mixed samples of DNA origami containing 1x and 2x, or 1x and 4x and 7x, target docking strands. Fitting results with the lowest error are highlighted in blue.

| | 1x+2x | | | | 1x+4x+7x | | | |
|---|---|---|---|---|---|---|---|---|
| | 2 peaks fitted (w1=w(1x)) (w2=w(2x)) | 2 peaks fitted (w1=w(1x)) (w2=w(3x)) | 1 peak fitted (w1=w(2x)) | 1 peak fitted (w1=w(1x)) | 3 peaks fitted (w1=w(1x)) (w2=w(4x)) (w3=w(7x)) | 3 peaks fitted (w1=w(1x)) (w2=w(3x)) (w3=w(7x)) | 2 peaks fitted (w1=w(1x)) (w2=w(5x)) | 2 peaks fitted (w1=w(1x)) (w2=w(7x)) |
| Chi-squared | 6E-6 | 2E-5 | 1E-5 | 6E-5 | 7E-6 | 8E-6 | 1E-5 | 1E-5 |
| R-squared | 0.995 | 0.986 | 0.986 | 0.957 | 0.981 | 0.980 | 0.958 | 0.965 |

**Table S6**: Fitting results for quantitative analysis of transfected EGFR in CHO cells. Fixed values are indicated in red. Errors are given as the standard error of the Gaussian fit.

| | Resting cells | | EGF-treated cells | | |
|---|---|---|---|---|---|
| | Peak 1 (background) | Peak 2 (signal monomer) | Peak 1 (background) | Peak 2 (signal monomer) | Peak 3 (signal dimer) |
| Mean / a. u. | 2.2 ± 0.1 | 7.0 ± 0.8 | 2.7 ± 0.1 | 6.4 ± 0.9 | 16 ± 3 |
| FWHM / a. u. | 2.0 ± 0.1 | 11.0 ± 1.7 | 2.7 ± 0.3 | 11.0 | 15.5 |
| Fraction / % | 41 ± 4 | 59 ± 10 | 27 ± 4 | 54 ± 5 | 20 ± 6 |

**Table S7**: Corrected EGFR dimer fractions. Calculations were based on the assumption that transfected cells only exhibit EGFR monomers and dimers. Errors are given as standard error.

| | Corrected dimer fraction / % |
|---|---|
| First peak fraction | 56 ± 8 |
| Second peak fraction | 52 ± 5 |
| Third peak fraction | 58 ± 17 |

**Table S8**: Cluster density of EGFR for transfected and untransfected cells. Errors are reported as standard errors.

| Cell condition | Nanobody | Cluster density / $\mu m^{-2}$ |
|---|---|---|
| transfected resting | NB@GFP-STAR 635P | 5.7 ± 1.3 |
| | NB@ALFA-4xP3 | 6.3 ± 1.3 |
| transfected EGF stimulated | NB@GFP-STAR 635P | 5.4 ± 1.5 |
| | NB@ALFA-4xP3 | 6 ± 3 |
| no transfection | NB@GFP-STAR 635P | 0.2 ± 0.1 |
| | NB@ALFA-4xP3 | 0.4 ± 0.3 |
| | NB@ALFA-4xP3 (no pre-incubation with imager strand) | 10 ± 3 |

**Table S9**: Staple strand sequences and strand extensions (extension sequences are given in **Table S1**) of the rectangular DNA origami.

| Posi-tion | Sequence (5' to 3'; binding to scaffold) | Strand extension |
|---|---|---|
| A1 | TTTTCACTCAAAGGGCGAAAAACCATCACC | |
| B1 | CAAATCAAGTTTTTTGGGGTCGAAACGTGGA | |
| C1 | AGCTGATTGCCCTTCAGAGTCCACTATTAAAGGGTGCCGT | |
| D1 | AAAGCACTAAATCGGAACCCTAATCCAGTT | |
| E1 | AGCAAGCGTAGGGTTGAGTGTTGTAGGGAGCC | |
| F1 | CCCGATTTAGAGCTTGACGGGGAAAAAGAATA | |
| G1 | CCCAGCAGGCGAAAAATCCCTTATAAATCAAGCCGGCG | |
| H1 | TCAATATCGAACCTCAAATATCAATTCCGAAA | |
| I1 | AACGTGGCGAGAAAGGAAGGGAAACCAGTAA | |
| J1 | TAAAAGGGACATTCTGGCCAACAAAGCATC | |
| K1 | TCAACAGTTGAAAGGAGCAAATGAAAAATCTAGAGATAGA | |
| L1 | ACCCTTCTGACCTGAAAGCGTAAGACGCTGAG | 2xP2 docking strand (applied for all) |
| M1 | CTTTAGGGCCTGCAACAGTGCCAATACGTG | 2xP2 docking strand (applied for all) |
| N1 | GCACAGACAATATTTTTGAATGGGGTCAGTA | 2xP2 docking strand (applied for all) |
| O1 | AGATTAGAGCCGTCAAAAAACAGAGGTGAGGCCTATTAGT | |
| P1 | CTTTAATGCGCGAACTGATAGCCCCACCAG | |
| A2 | GTCGACTTCGGCCAACGCGCGGGGTTTTTC | |
| B2 | CTCCAACGCAGTGAGACGGGCAACCAGCTGCA | |
| D2 | TGGAACAACCGCCTGGCCCTGAGGCCCGCT | |
| E2 | CTGTGTGATTGCGTTGCGCTCACTAGAGTTGC | |
| F2 | GCCCGAGAGTCCACGCTGGTTTGCAGCTAACT | |
| H2 | GCAATTCACATATTCCTGATTATCAAAGTGTA | |
| I2 | TCGGCAAATCCTGTTTGATGGTGGACCCTCAA | |
| J2 | ACCTTGCTTGGTCAGTTGGCAAAGAGCGGA | |
| L2 | AGCCAGCAATTGAGGAAGGTTATCATCATTTT | 2xP2 docking strand (applied for all) |
| M2 | CTACCATAGTTTGAGTAACATTTAAAATAT | 2xP2 docking strand (applied for all) |
| N2 | TTAACACCAGCACTAACAACTAATCGTTATTA | 2xP2 docking strand (applied for all) |
| P2 | CAGAAGATTAGATAATACATTTGTCGACAA | |
| A3 | TGCATCTTTCCCAGTCACGACGGCCTGCAG | |
| B3 | TTAATGAACTAGAGGATCCCCGGGGGGTAACG | |
| D3 | TTCCAGTCGTAATCATGGTCATAAAAGGGG | |
| E3 | GCTTTCCGATTACGCCAGCTGGCGGCTGTTTC | |
| F3 | CACATTAAAATTGTTATCCGCTCATGCGGGCC | |
| H3 | AGAAAACAAAGAAGATGATGAAACAGGCTGCG | |
| I3 | AAGCCTGGTACGAGCCGGAAGCATAGATGATG | |
| J3 | ATTATCATTCAATATAATCCTGACAATTAC | |
| L3 | GCGGAACATCTGAATAATGGAAGGTACAAAAT | |
| M3 | CATAAATCTTTGAATACCAAGTGTTAGAAC | 2xP2 docking strand (applied for all) |
| N3 | ATTTTAAAATCAAAATTATTTGCACGGATTCG | |
| P3 | CTCGTATTAGAAATTGCGTAGATACAGTAC | |
| A4 | TAATCAGCGGATTGACCGTAATCGTAACCG | |
| B4 | CCAGGGTTGCCAGTTTGAGGGGACCCGTGGGA | |
| C4 | GTATAAGCCAACCCGTCGGATTCTGACGACAGTATCGGCCGCAAGGCG | |
| D4 | GATGTGCTTCAGGAAGATCGCACAATGTGA | |
| E4 | ATATTTTGGCTTTCATCAACATTATCCAGCCA | |
| F4 | TCTTCGCTGCACCGCTTCTGGTGCGGCCTTCC | |
| G4 | TAAATCAAAATAATTCGCGTCTCGGAAACCAGGCAAAGGGAAGG | |
| H4 | ATCGCAAGTATGTAAATGCTGATGATAGGAAC | |
| I4 | CAACTGTTGCGCCATTCGCCATTCAAACATCA | |
| J4 | CTGAGCAAAAATTAATTACATTTTGGGTTA | |

| | | |
|---|---|---|
| K4 | TCAAATATAACCTCCGGCTTAGGTAACAATTTCATTTGAAGGCGAATT | |
| L4 | CGCGCAGATTACCTTTTTTAATGGGAGAGACT | |
| M4 | CCTAAATCAAAATCATAGGTCTAAACAGTA | |
| N4 | CCTGATTGCAATATATGTGAGTGATCAATAGT | |
| O4 | GTGATAAAAAGACGCTGAGAAGAGATAACCTTGCTTCTGTTCGGGAGA | |
| P4 | CTTTTACAAAATCGTCGCTATTAGCGATAG | |
| A5 | AACGCAAAATCGATGAACGGTACCGGTTGA | |
| B5 | ACAAACGGAAAAGCCCCAAAAACACTGGAGCA | |
| C5 | TATATTTTGTCATTGCCTGAGAGTGGAAGATT | |
| D5 | GCGAGTAAAAATATTTAAATTGTTACAAAG | |
| E5 | TAGGTAAACTATTTTTGAGAGATCAAACGTTA | |
| F5 | TGTAGCCATTAAAATTCGCATTAAATGCCGGA | |
| G5 | GAGACAGCTAGCTGATAAATTAATTTTTGT | |
| H5 | GTAATAAGTTAGGCAGAGGCATTTATGATATT | |
| I5 | GCCATCAAGCTCATTTTTTAACCACAAATCCA | |
| J5 | TATAACTAACAAAGAACGCGAGAACGCCAA | |
| K5 | GTAAAGTAATCGCCATATTTAACAAAACTTTT | |
| L5 | ACCTTTTTATTTTAGTTAATTTCATAGGGCTT | |
| M5 | ACAACATGCCAACGCTCAACAGTCTTCTGA | |
| N5 | GAATTTATTTAATGGTTTGAAATATTCTTACC | |
| O5 | GTTTATCAATATGCGTTATACAAACCGACCGT | |
| P5 | CTTAGATTTAAGGCGTTAAATAAAGCCTGT | |
| A6 | AACAGTTTTGTACCAAAAACATTTTATTTC | 2xP2 docking strand (applied for all) |
| B6 | AACAAGAGGGATAAAAATTTTTAGCATAAAGC | |
| C6 | GATTTAGTCAATAAAGCCTCAGAGAACCCTCA | |
| D6 | GCTATCAGAAATGCAATGCCTGAATTAGCA | |
| E6 | AATGGTCAACAGGCAAGGCAAAGAGTAATGTG | |
| F6 | GAGGGTAGGATTCAAAAGGGTGAGACATCCAA | |
| G6 | TTTGGGGATAGTAGTAGCATTAAAAGGCCG | |
| H6 | CCAATAGCTCATCGTAGGAATCATGGCATCAA | 4xP3 docking strand (applied for 7x) |
| I6 | CAACCGTTTCAAATCACCATCAATTCGAGCCA | 4xP3 docking strand (applied for 3x-5x,7x) |
| J6 | CATGTAATAGAATATAAAGTACCAAGCCGT | 4xP3 docking strand (applied for 3x-5x,7x) |
| K6 | TATCCGGTCTCATCGAGAACAAGCGACAAAAG | |
| L6 | AATTGAGAATTCTGTCCAGACGACTAAACCAA | |
| M6 | GCGAACCTCCAAGAACGGGTATGACAATAA | |
| N6 | AGTATAAAGTTCAGCTAATGCAGATGTCTTTC | |
| O6 | GCCTTAAACCAATCAATAATCGGCACGCGCCT | |
| P6 | TTAGTATCACAATAGATAAGTCCACGAGCA | |
| A7 | TTTACCCCAACATGTTTTAAATTTCCATAT | 2xP2 docking strand (applied for all) |
| B7 | TAAATCGGGATTCCCAATTCTGCGATATAATG | 2xP2 docking strand (applied for all) |
| C7 | CGGATTGCAGAGCTTAATTGCTGAAACGAGTA | 2xP2 docking strand (applied for all) |
| D7 | AAATTAAGTTGACCATTAGATACTTTTGCG | |
| E7 | CGAAAGACTTTGATAAGAGGTCATATTTCGCA | |
| F7 | TAAATCATATAACCTGTTTAGCTAACCTTTAA | |
| G7 | GCTTCAATCAGGATTAGAGAGTTATTTTCA | |
| H7 | AGAGAGAAAAAAATGAAAATAGCAAGCAAACT | 4xP3 docking strand (applied for 4x,5x,7x) |
| I7 | TTCTACTACGCGAGCTGAAAAGGTTACCGCGC | 4xP3 docking strand (applied for 1x-5x,7x) |
| J7 | TTTTATTTAAGCAAATCAGATATTTTTTGT | 4xP3 docking strand (applied for 7x) |
| K7 | TTAGACGGCCAAATAAGAAACGATAGAAGGCT | |
| L7 | GTACCGCAATTCTAAGAACGCGAGTATTATTT | |
| M7 | AAAGTCACAAAATAAACAGCCAGCGTTTTA | |
| N7 | CTTATCATTCCCGACTTGCGGGAGCCTAATTT | |
| O7 | GAGAGATAGAGCGTCTTTCCAGAGGTTTTGAA | |

| | | |
|---|---|---|
| P7 | TGTAGAAATCAAGATTAGTTGCTCTTACCA | |
| A8 | TTTAGGACAAATGCTTTAAACAATCAGGTC | 2xP2 docking strand (applied for all) |
| B8 | CTGTAGCTTGACTATTATAGTCAGTTCATTGA | 2xP2 docking strand (applied for all) |
| C8 | ATGCAGATACATAACGGGAATCGTCATAAATAAAGCAAAG | 2xP2 docking strand (applied for all) |
| D8 | GATGGCTTATCAAAAAGATTAAGAGCGTCC | |
| E8 | TAAGAGCAAATGTTTAGACTGGATAGGAAGCC | |
| F8 | TTGCTCCTTTCAAATATCGCGTTTGAGGGGGT | |
| G8 | CGTTTACCAGACGACAAAGAAGTTTTGCCATAATTCGA | |
| H8 | TTATTACGAAGAACTGGCATGATTGCGAGAGG | |
| I8 | CCAACAGGAGCGAACCAGACCGGAGCCTTTAC | 4xP3 docking strand (applied for 5x,7x) |
| J8 | TTAACGTCTAACATAAAAACAGGTAACGGA | |
| K8 | CGTAGAAAATACATACCGAGGAAACGCAATAAGAAGCGCA | |
| L8 | ATCCCAATGAGAATTAACTGAACAGTTACCAG | |
| M8 | AACGCAAAGATAGCCGAACAAACCCTGAAC | |
| N8 | GCCAGTTAGAGGGTAATTGAGCGCTTTAAGAA | |
| O8 | GTTTATTTTGTCACAATCTTACCGAAGCCCTTTAATATCA | |
| P8 | ACGCTAACACCCACAAGAATTGAAAATAGC | |
| A9 | CATCAAGTAAAACGAACTAACGAGTTGAGA | |
| B9 | ATCCCCCTATACCACATTCAACTAGAAAAATC | |
| D9 | AATACTGCCCAAAAGGAATTACGTGGCTCA | |
| E9 | TCATTCAGATGCGATTTTAAGAACAGGCATAG | |
| F9 | AATAGTAAACACTATCATAACCCTCATTGTGA | |
| H9 | GCAAGGCCTCACCAGTAGCACCATGGGCTTGA | |
| I9 | CTTTTGCAGATAAAAACCAAAATAAAGACTCC | |
| J9 | ATACCCAACAGTATGTTAGCAAATTAGAGC | |
| L9 | AAGGAAACATAAAGGTGGCAACATTATCACCG | |
| M9 | TCAAGTTTCATTAAAGGTGAATATAAAAGA | |
| N9 | AAGTAAGCAGACACCACGGAATAATATTGACG | |
| P9 | AATAGCTATCAATAGAAAATTCAACATTCA | |
| A10 | AATACGTTTGAAAGAGGACAGACTGACCTT | |
| B10 | TACGTTAAAGTAATCTTGACAAGAACCGAACT | |
| D10 | TTATACCACCAAATCAACGTAACGAACGAG | |
| E10 | ACACTCATCCATGTTACTTAGCCGAAAGCTGC | |
| F10 | ATTACCTTTGAATAAGGCTTGCCCAAATCCGC | |
| H10 | TTGACAGGCCACCACCAGAGCCGCGATTTGTA | |
| I10 | GATGGTTTGAACGAGTAGTAAATTTACCATTA | |
| J10 | CAGCAAAAGGAAACGTCACCAATGAGCCGC | |
| L10 | TCACCGACGCACCGTAATCAGTAGCAGAACCG | 2xP2 docking strand (applied for all) |
| M10 | TTAAAGCCAGAGCCGCCACCCTCGACAGAA | 2xP2 docking strand (applied for all) |
| N10 | GAAATTATTGCCTTTAGCGTCAGACCGGAACC | 2xP2 docking strand (applied for all) |
| P10 | ACCGATTGTCGGCATTTTCGGTCATAATCA | |
| A11 | AGGCTCCAGAGGCTTTGAGGACACGGGTAA | |
| B11 | GACCAACTAATGCCACTACGAAGGGGGTAGCA | |
| C11 | TTTATCAGGACAGCATCGGAACGACACCAACCTAAAACGAGGTCAATC | |
| D11 | GCGCAGACAAGAGGCAAAAGAATCCCTCAG | |
| E11 | AAACAGCTTTTTGCGGGATCGTCAACACTAAA | |
| F11 | GACCTGCTCTTTGACCCCCAGCGAGGGAGTTA | |
| G11 | TGACAACTCGCTGAGGCTTGCATTATACCAAGCGCGATGATAAA | |
| H11 | TTAGGATTGGCTGAGACTCCTCAATAACCGAT | |
| I11 | TCATCGCCAACAAAGTACAACGGACGCCAGCA | |
| J11 | CACCAGAAAGGTTGAGGCAGGTCATGAAAG | |
| K11 | GCGGATAACCTATTATTCTGAAACAGACGATTGGCCTTGAAGAGCCAC | |
| L11 | CCACCCTCTATTCACAAACAAATACCTGCCTA | 2xP2 docking strand (applied for all) |

| | | |
|---|---|---|
| M11 | GTATAGCAAACAGTTAATGCCCAATCCTCA | 2xP2 docking strand (applied for all) |
| N11 | GCCTCCCTCAGAATGGAAAGCGCAGTAACAGT | 2xP2 docking strand (applied for all) |
| O11 | CAGGAGGTGGGGTCAGTGCCTTGAGTCTCTGAATTTACCGGGAACCAG | |
| P11 | AAATCACCTTCCAGTAAGCGTCAGTAATAA | |
| A12 | AGAAAGGAACAACTAAAGGAATTCAAAAAAA | |
| B12 | ACGGCTACAAAAGGAGCCTTTAATGTGAGAAT | |
| C12 | ACAACTTTCAACAGTTTCAGCGGATGTATCGG | |
| D12 | CAGCGAAACTTGCTTTCGAGGTGTTGCTAA | |
| E12 | TAAATGAATTTTCTGTATGGGATTAATTTCTT | |
| F12 | AAGGCCGCTGATACCGATAGTTGCGACGTTAG | |
| G12 | TCTAAAGTTTTGTCGTCTTTCCAGCCGACAA | |
| H12 | TCCACAGACAGCCCTCATAGTTAGCGTAACGA | |
| I12 | ATATTCGGAACCATCGCCCACGCAGAGAAGGA | |
| J12 | TATTAAGAAGCGGGGTTTTGCTCGTAGCAT | |
| K12 | TCACCAGTACAAACTACAACGCCTAGTACCAG | |
| L12 | TTTCGGAAGTGCCGTCGAGAGGGTGAGTTTCG | |
| M12 | AGGAACCCATGTACCGTAACACTTGATATAA | 2xP2 docking strand (applied for all) |
| N12 | GCCCGTATCCGGAATAGGTGTATCAGCCCAAT | |
| O12 | CCACCCTCATTTTCAGGGATAGCAACCGTACT | |
| P12 | GTTTTAACTTAGTACCGCCACCCAGAGCCA | |

**Table S10:** Sequences of biotinylated staple strands for DNA origami.

| **Position** | **Sequence** | **Modification** |
|---|---|---|
| C2 | ATTAAGTTTACCGAGCTCGAATTCGGGAAACCTGTCGTGC | 5'-biotin |
| C9 | ATAAGGGAACCGGATATTCATTACGTCAGGACGTTGGGAA | 5'-biotin |
| G2 | GCGATCGGCAATTCCACACAACAGGTGCCTAATGAGTG | 5'-biotin |
| G9 | TTGTGTCGTGACGAGAAACACCAAATTTCAACTTTAAT | 5'-biotin |
| K2 | ATTCATTTTTGTTTGGATTATACTAAGAAACCACCAGAAG | 5'-biotin |
| K9 | CACCCTCAGAAACCATCGATAGCATTGAGCCATTTGGGAA | 5'-biotin |
| O2 | AACAATAACGTAAAACAGAAATAAAAATCCTTTGCCCGAA | 5'-biotin |
| O9 | AGCCACCACTGTAGCGCGTTTTCAAGGGAGGGAAGGTAAA | 5'-biotin |

**Table S11:** Sequence of scaffold strand (M13mp18) for DNA origami.

AATGCTACTACTATTAGTAGAATTGATGCCACCTTTTCAGCTCGCGCCCCAAATGAAAATATAGCTAAACAGGTTATTGACCATTTGCGAAATGTAT
CTAATGGTCAAACTAAATCTACTCGTTCGCAGAATTGGGAATCAACTGTTATATGGAATGAAACTTCCAGACACCGTACTTTAGTTGCATATTTAAA
ACATGTTGAGCTACAGCATTATATTCAGCAATTAAGCTCTAAGCCATCCGCAAAAATGACCTCTTATCAAAAGGAGCAATTAAAGGTACTCTCTAA
TCCTGACCTGTTGGAGTTTGCTTCCGGTCTGGTTCGCTTTGAAGCTCGAATTAAAACGCGATATTTGAAGTCTTTCGGGCTTCCTCTTAATCTTT
TTGATGCAATCCGCTTTGCTTCTGACTATAATAGTCAGGGTAAAGACCTGATTTTTGATTTATGGTCATTCTCGTTTTCTGAACTGTTTAAAGCATT
TGAGGGGGATTCAATGAATATTTATGACGATTCCGCAGTATTGGACGCTATCCAGTCTAAACATTTTACTATTACCCCCTCTGGCAAAACTTCTTT
TGCAAAAGCCTCTCGCTATTTTGGTTTTTATCGTCGTCTGGTAAACGAGGGTTATGATAGTGTTGCTCTTACTATGCCTCGTAATTCCTTTTGGCG
TTATGTATCTGCATTAGTTGAATGTGGTATTCCTAAATCTCAACTGATGAATCTTTCTACCTGTAATAATGTTGTTCCGTTAGTTCGTTTTATTAACG
TAGATTTTTCTTCCCAACGTCCTGACTGGTATAATGAGCCAGTTCTTAAAATCGCATAAGGTAATTCACAATGATTAAAGTTGAAATTAAACCATCT
CAAGCCCAATTTACTACTCGTTCTGGTGTTTCTCGTCAGGGCAAGCCTTATTCACTGAATGAGCAGCTTTGTTACGTTGATTTGGGTAATGAATA
TCCGGTTCTTGTCAAGATTACTCTTGATGAAGGTCAGCCAGCCTATGCGCCTGGTCTGTACACCGTTCATCTGTCCTCTTTCAAAGTTGGTCAG
TTCGGTTCCCTTATGATTGACCGTCTGCGCCTCGTTCCGGCTAAGTAACATGGAGCAGGTCGCGGATTTCGACACAATTTATCAGGCGATGATA
CAAATCTCCGTTGTACTTTGTTTCGCGCTTGGTATAATCGCTGGGGGTCAAAGATGAGTGTTTTAGTGTATTCTTTTGCCTCTTTCGTTTTAGGTT
GGTGCCTTCGTAGTGGCATTACGTATTTTACCCGTTTAATGGAAACTTCCTCATGAAAAAGTCTTTAGTCCTCAAAGCCTCTGTAGCCGTTGCTA
CCCTCGTTCCGATGCTGTCTTTCGCTGCTGAGGGTGACGATCCCGCAAAAGCGGCCTTTAACTCCCTGCAAGCCTCAGCGACCGAATATATCG
GTTATGCGTGGGCGATGGTTGTTGTCATTGTCGGCGCAACTATCGGTATCAAGCTGTTTAAGAAATTCACCTCGAAAGCAAGCTGATAAACCGA
TACAATTAAAGGCTCCTTTTGGAGCCTTTTTTTTGGAGATTTTCAACGTGAAAAAATTATTATTCGCAATTCCTTTAGTTGTTCCTTTCTATTCTCAC
TCCGCTGAAACTGTTGAAAGTTGTTTAGCAAAATCCCATACAGAAAATTCATTTACTAACGTCTGGAAAGACGACAAAACTTTAGATCGTTACGC
TAACTATGAGGGCTGTCTGTGGAATGCTACAGGCGTTGTAGTTTGTACTGGTGACGAAACTCAGTGTTACGGTACATGGGTTCCTATTGGGCTT
GCTATCCCTGAAAATGAGGGTGGTGGCTCTGAGGGTGGCGGTTCTGAGGGTGGCGGTTCTGAGGGTGGCGGTACTAAACCTCCTGAGTACG
GTGATACACCTATTCCGGGCTATACTTATATCAACCCTCTCGACGGCACTTATCCGCCTGGTACTGAGCAAAACCCCGCTAATCCTAATCCTTCT
CTTGAGGAGTCTCAGCCTCTTAATACTTTCATGTTTCAGAATAATAGGTTCCGAAATAGGCAGGGGGCATTAACTGTTTATACGGGCACTGTTAC
TCAAGGCACTGACCCCGTTAAAACTTATTACCAGTACACTCCTGTATCATCAAAAGCCATGTATGACGCTTACTGGAACGGTAAATTCAGAGACT
GCGCTTTCCATTCTGGCTTTAATGAGGATTTATTTGTTTGTGAATATCAAGGCCAATCGTCTGACCTGCCTCAACCTCCTGTCAATGCTGGCGGC
GGCTCTGGTGGTGGTTCTGGTGGCGGCTCTGAGGGTGGTGGCTCTGAGGGTGGCGGTTCTGAGGGTGGCGGCTCTGAGGGAGGCGGTTC
CGGTGGTGGCTCTGGTTCCGGTGATTTTGATTATGAAAAGATGGCAAACGCTAATAAGGGGGCTATGACCGAAAATGCCGATGAAAACGCGCT
ACAGTCTGACGCTAAAGGCAAACTTGATTCTGTCGCTACTGATTACGGTGCTGCTATCGATGGTTTCATTGGTGACGTTTCCGGCCTTGCTAAT
GGTAATGGTGCTACTGGTGATTTTGCTGGCTCTAATTCCCAAATGGCTCAAGTCGGTGACGGTGATAATTCACCTTTAATGAATAATTTCCGTCA
ATATTTACCTTCCCTCCCTCAATCGGTTGAATGTCGCCCTTTTGTCTTTGGCGCTGGTAAACCATATGAATTTTCTATTGATTGTGACAAAATAAAC
TTATTCCGTGGTGTCTTTGCGTTTCTTTTATATGTTGCCACCTTTATGTATGTATTTTCTACGTTTGCTAACATACTGCGTAATAAGGAGTCTTAATC
ATGCCAGTTCTTTTGGGTATTCCGTTATTATTGCGTTTCCTCGGTTTCCTTCTGGTAACTTTGTTCGGCTATCTGCTTACTTTTCTTAAAAAGGGC
TTCGGTAAGATAGCTATTGCTATTTCATTGTTTCTTGCTCTTATTATTGGGCTTAACTCAATTCTTGTGGGTTATCTCTCTGATATTAGCGCTCAATT
ACCCTCTGACTTTGTTCAGGGTGTTCAGTTAATTCTCCCGTCTAATGCGCTTCCCTGTTTTTATGTTATTCTCTCTGTAAAGGCTGCTATTTTCATT

TTTGACGTTAAACAAAAAATCGTTTCTTATTTGGATTGGGATAAATAATATGGCTGTTTATTTTGTAACTGGCAAATTAGGCTCTGGAAAGACGCTC
GTTAGCGTTGGTAAGATTCAGGATAAAATTGTAGCTGGGTGCAAAATAGCAACTAATCTTGATTTAAGGCTTCAAAACCTCCCGCAAGTCGGGA
GGTTCGCTAAAACGCCTCGCGTTCTTAGAATACCGGATAAGCCTTCTATATCTGATTTGCTTGCTATTGGGCGCGGTAATGATTCCTACGATGAA
AATAAAAACGGCTTGCTTGTTCTCGATGAGTGCGGTACTTGGTTTAATACCCGTTCTTGGAATGATAAGGAAAGACAGCCGATTATTGATTGGTT
TCTACATGCTCGTAAATTAGGATGGGATATTATTTTTCTTGTTCAGGACTTATCTATTGTTGATAAACAGGCGCGTTCTGCATTAGCTGAACATGTT
GTTTATTGTCGTCGTCTGGACAGAATTACTTTACCTTTTGTCGGTACTTTATATTCTCTTATTACTGGCTCGAAAATGCCTCTGCCTAAATTACATG
TTGGCGTTGTTAAATATGGCGATTCTCAATTAAGCCCTACTGTTGAGCGTTGGCTTTATACTGGTAAGAATTTGTATAACGCATATGATACTAAACA
GGCTTTTTCTAGTAATTATGATTCCGGTGTTTATTCTTATTTAACGCCTTATTTATCACACGGTCGGTATTTCAAACCATTAAATTTAGGTCAGAAGA
TGAAATTAACTAAAATATATTTGAAAAAGTTTTCTCGCGTTCTTTGTCTTGCGATTGGATTTGCATCAGCATTTACATATAGTTATATAACCCAACCT
AAGCCGGAGGTTAAAAAGGTAGTCTCTCAGACCTATGATTTTGATAAATTCACTATTGACTCTTCTCAGCGTCTTAATCTAAGCTATCGCTATGTTT
TCAAGGATTCTAAGGGAAAATTAATTAATAGCGACGATTTACAGAAGCAAGGTTATTCACTCACATATATTGATTTATGTACTGTTTCCATTAAAAAA
GGTAATTCAAATGAAATTGTTAAATGTAATTAATTTTGTTTTCTTGATGTTTGTTTCATCATCTTCTTTTGCTCAGGTAATTGAAATGAATAATTCGCC
TCTGCGCGATTTTGTAACTTGGTATTCAAAGCAATCAGGCGAATCCGTTATTGTTTCTCCCGATGTAAAAGGTACTGTTACTGTATATTCATCTGA
CGTTAAACCTGAAAATCTACGCAATTTCTTTATTTCTGTTTTACGTGCAAATAATTTTGATATGGTAGGTTCTAACCCTTCCATTATTCAGAAGTATA
ATCCAAACAATCAGGATTATATTGATGAATTGCCATCATCTGATAATCAGGAATATGATGATAATTCCGCTCCTTCTGGTGGTTTCTTTGTTCCGCA
AAATGATAATGTTACTCAAACTTTTAAAATTAATAACGTTCGGGCAAAGGATTTAATACGAGTTGTCGAATTGTTTGTAAAGTCTAATACTTCTAAAT
CCTCAAATGTATTATCTATTGACGGCTCTAATCTATTAGTTGTTAGTGCTCCTAAAGATATTTTAGATAACCTTCCTCAATTCCTTTCAACTGTTGAT
TTGCCAACTGACCAGATATTGATTGAGGGTTTGATATTTGAGGTTCAGCAAGGTGATGCTTTAGATTTTTCATTTGCTGCTGGCTCTCAGCGTGG
CACTGTTGCAGGCGGTGTTAATACTGACCGCCTCACCTCTGTTTTATCTTCTGCTGGTGGTTCGTTCGGTATTTTTAATGGCGATGTTTTAGGGC
TATCAGTTCGCGCATTAAAGACTAATAGCCATTCAAAAATATTGTCTGTGCCACGTATTCTTACGCTTTCAGGTCAGAAGGGTTCTATCTCTGTTG
GCCAGAATGTCCCTTTTATTACTGGTCGTGTGACTGGTGAATCTGCCAATGTAAATAATCCATTTCAGACGATTGAGCGTCAAAATGTAGGTATTT
CCATGAGCGTTTTTCCTGTTGCAATGGCTGGCGGTAATATTGTTCTGGATATTACCAGCAAGGCCGATAGTTTGAGTTCTTCTACTCAGGCAAGT
GATGTTATTACTAATCAAAGAAGTATTGCTACAACGGTTAATTTGCGTGATGGACAGACTCTTTTACTCGGTGGCCTCACTGATTATAAAAACACT
TCTCAGGATTCTGGCGTACCGTTCCTGTCTAAAATCCCTTTAATCGGCCTCCTGTTTAGCTCCCGCTCTGATTCTAACGAGGAAAGCACGTTATA
CGTGCTCGTCAAAGCAACCATAGTACGCGCCCTGTAGCGGCGCATTAAGCGCGGCGGGTGTGGTGGTTACGCGCAGCGTGACCGCTACACT
TGCCAGCGCCCTAGCGCCCGCTCCTTTCGCTTTCTTCCCTTCCTTTCTCGCCACGTTCGCCGGCTTTCCCCGTCAAGCTCTAAATCGGGGGC
TCCCTTTAGGGTTCCGATTTAGTGCTTTACGGCACCTCGACCCCAAAAAACTTGATTTGGGTGATGGTTCACGTAGTGGGCCATCGCCCTGATA
GACGGTTTTTCGCCCTTTGACGTTGGAGTCCACGTTCTTTAATAGTGGACTCTTGTTCCAAACTGGAACAACACTCAACCCTATCTCGGGCTAT
TCTTTTGATTTATAAGGGATTTTGCCGATTTCGGAACCACCATCAAACAGGATTTTCGCCTGCTGGGGCAAACCAGCGTGGACCGCTTGCTGCA
ACTCTCTCAGGGCCAGGCGGTGAAGGGCAATCAGCTGTTGCCCGTCTCACTGGTGAAAAGAAAAACCACCCTGGCGCCCAATACGCAAACC
GCCTCTCCCCGCGCGTTGGCCGATTCATTAATGCAGCTGGCACGACAGGTTTCCCGACTGGAAAGCGGGCAGTGAGCGCAACGCAATTAATG
TGAGTTAGCTCACTCATTAGGCACCCCAGGCTTTACACTTTATGCTTCCGGCTCGTATGTTGTGTGGAATTGTGAGCGGATAACAATTTCACACA
GGAAACAGCTATGACCATGATTACGAATTCGAGCTCGGTACCCGGGGATCCTCTAGAGTCGACCTGCAGGCATGCAAGCTTGGCACTGGCCG
TCGTTTTACAACGTCGTGACTGGGAAAACCCTGGCGTTACCCAACTTAATCGCCTTGCAGCACATCCCCCTTTCGCCAGCTGGCGTAATAGCG
AAGAGGCCCGCACCGATCGCCCTTCCCAACAGTTGCGCAGCCTGAATGGCGAATGGCGCTTTGCCTGGTTTCCGGCACCAGAAGCGGTGCC
GGAAAGCTGGCTGGAGTGCGATCTTCCTGAGGCCGATACTGTCGTCGTCCCCTCAAACTGGCAGATGCACGGTTACGATGCGCCCATCTACA
CCAACGTGACCTATCCCATTACGGTCAATCCGCCGTTTGTTCCCACGGAGAATCCGACGGGTTGTTACTCGCTCACATTTAATGTTGATGAAAG
CTGGCTACAGGAAGGCCAGACGCGAATTATTTTTGATGGCGTTCCTATTGGTTAAAAAATGAGCTGATTTAACAAAAATTTAATGCGAATTTTAAC
AAAATATTAACGTTTACAATTTAAATATTTGCTTATACAATCTTCCTGTTTTTGGGGCTTTTCTGATTATCAACCGGGGTACATATGATTGACATGCTA
GTTTTACGATTACCGTTCATCGATTCTCTTGTTTGCTCCAGACTCTCAGGCAATGACCTGATAGCCTTTGTAGATCTCTCAAAAATAGCTACCCTC
TCCGGCATTAATTTATCAGCTAGAACGGTTGAATATCATATTGATGGTGATTTGACTGTCTCCGGCCTTTCTCACCCTTTTGAATCTTTACCTACAC
ATTACTCAGGCATTGCATTTAAAATATATGAGGGTTCTAAAAATTTTTATCCTTGCGTTGAAATAAAGGCTTCTCCCGCAAAAGTATTACAGGGTCA
TAATGTTTTTGGTACAACCGATTTAGCTTTATGCTCTGAGGCTTTATTGCTTAATTTTGCTAATTCTTTGCCTTGCCTGTATGATTTATTGGATGTT

**Table S12**: Imaging settings in microscopy experiments shown in main and supplemental figures.

| Figure | 571-630 nm detection | | 650-763 nm detection | | STED PSF modulation | Pinhole / AU | Line accumulations | Pixel dwell time / µs | Pixel size / nm |
|---|---|---|---|---|---|---|---|---|---|
| | 561 nm laser power / µW | 775 nm laser power / mW | 640 nm laser power / µW | 775 nm laser power / mW | | | | | |
| 1D, S1 | 5.7 | 240 | 4.7 | 240 | 2D | 1.0 | 20 | 5 | 12,5 |
| 2, 3 | 3.7 | 157 | 3.2 | 109 | 2D | 1.0 | 20-25 | 5 | 30 |
| 4, S5 (confocal) | 3.3 | - | 3.2 | - | - | 1.0 | 5-10 | 5 | 50 |
| 4, S5 (STED) | 3.3 | 109 | 3.2 | 91 | 2D | 0.81 | 20-35 | 5 | 30 |
| S6 | 1.5 | 45 | 3.2 | 28 | 3D (top hat) | 0.71 | 7 | 17 | 60 (voxel size) |

**Table S13**: Fraction of specific DNA origami signal analyzed from the total identified DNA Origami signal. Errors are given as standard errors.

| | 1x | 2x | 3x | 4x | 5x | 7x |
|---|---|---|---|---|---|---|
| Fraction of specific signal / % | 92.1 ± 1.1 | 97.0 ± 1.4 | 94 ± 3 | 96.9 ± 1.7 | 96.3 ± 1.8 | 97 ± 3 |
